# Extracellular DNA controls expression of *Pseudomonas aeruginosa* genes involved in nutrient utilization, metal homeostasis, acid pH tolerance, and virulence

**DOI:** 10.1101/850446

**Authors:** Shawn Lewenza, Lori Johnson, Laetitia Charron-Mazenod, Mia Hong, Heidi Mulcahy-O’Grady

## Abstract

*Pseudomonas aeruginosa* grows in extracellular DNA-enriched biofilms and infection sites. Extracellular DNA (eDNA) is generally considered a structural biofilm polymer required for aggregation and biofilm maturation. In addition, eDNA can sequester divalent metal cations, acidify the growth media, and serve as a nutrient source. Here we determine the transcriptome of planktonic *P. aeruginosa* grown in the presence of eDNA using RNA-seq. Transcriptome analysis identified 89 induced genes and 76 repressed genes in the presence of eDNA (FDR<0.05), and transcriptional *lux* fusions were used to confirm eDNA regulation. A large number of eDNA-induced genes appear to be involved in utilizing DNA as a nutrient. Several eDNA-induced genes are also induced by acidic pH 5.5, and growth in the presence of eDNA or acidic pH promoted an acid tolerance response in *P. aeruginosa*. The *cyoABCDE* terminal oxidase is induced at pH 5.5 and contributed to the acid tolerance phenotype. Quantitative metal analysis confirmed that DNA binds to diverse metals, which helps explain why many genes involved in a general uptake of metals were controlled by eDNA. Growth in the presence of eDNA also promoted intracellular bacterial survival and influenced virulence and during the acute infection model of fruit flies. The diverse functions of the eDNA-regulated genes underscore the important role of this extracellular polymer in promoting antibiotic resistance, virulence, acid tolerance, and nutrient utilization; phenotypes that contribute to long-term survival.

## INTRODUCTION

Bacteria encounter the presence of extracellular DNA (eDNA) when growing as biofilms and during interactions with immune cells (1). During biofilm formation, eDNA in the extracellular matrix arises through autolysis, secretion, outer membrane vesicle (OMV) release or phage-mediated lysis (2). During infections, bacteria are likely exposed to eDNA in many infection sites, where microbes encounter neutrophil extracellular traps (NETs) (3). NETs are an ejected lattice of chromosomal DNA that is enmeshed with numerous antimicrobial proteins from neutrophil granules that function to trap and kill numerous microbial organisms (3). *P. aeruginosa* encounters NETs in the Cystic Fibrosis (CF) sputum (4,5) and also during eye and skin infections (6,3).

We are beginning to appreciate the functions of extracellular DNA on bacterial physiology through understanding its role in bacterial biofilms. Extracellular DNA is a ubiquitous biofilm matrix polymer that has been shown to promote attachment and biofilm formation in most bacterial species tested (7). Treatment of biofilms with deoxyribonuclease is a successful means to dissolve pre-formed biofilms and to prevent biofilm formation, and thus constitutes a novel biofilm treatment strategy (7). Biofilm matrix DNA can also influence the surface charge and bacterial adhesion, as well as influence the interactions with bacterial surface adhesins (2). Extracellular DNA coordinates the migration of *P. aeruginosa* aggregates in interstitial biofilms formed on semisolid media (8).

In previous studies, we explored the hypothesis that extracellular DNA influences bacterial gene expression. We demonstrated that DNA is a polyanion and has a large capacity to bind and sequester divalent cations (9). The addition of eDNA to bacterial cultures sequesters Mg^2+^ and therefore activates the PhoPQ and PmrAB two component systems, which both respond to limiting Mg^2+^ (1,9,10). At high concentrations, the cation chelating activity of DNA can disrupt the inner and outer membrane integrity, leading to rapid lysis and further DNA release (9).

The accumulation of extracellular DNA also acidifies planktonic and biofilm cultures, likely through protons donated by the phosphate groups along the DNA backbone (11). The accumulation of DNA in the biofilm matrix contributes to the pH gradients in biofilms, and may therefore contribute to the acidic pH of the CF lung environment (11). Both cation chelation and acidification are independent signals caused by extracellular DNA that induce expression of the PhoPQ/PmrAB-regulated genes. In these examples, the physicochemical properties of extracellular DNA shape the environmental conditions encountered by bacteria in DNA rich niches.

Several groups have shown that DNA is an efficient nutrient source of carbon, nitrogen and phosphate (12–14). Extracellular DNA and the DNA within neutrophil NETS induces the expression of a 2 gene operon encoding a secreted phosphatase and deoxyribonuclease in *P. aerugino*sa (14,15). These secreted enzymes target DNA and protect *P. aeruginosa* from DNA killing during exposure to neutrophil extracellular traps (15). These genes are also induced under phosphate limiting conditions, and also contribute to DNA degradation and phosphate acquisition when using DNA as a nutrient (14).

The focus of this study was to determine the global effect of extracellular DNA on the transcriptome of *P. aeruginosa*. We used RNA-seq to identify genes regulated by eDNA and provide evidence that eDNA promotes an acid tolerance response, influences *P. aeruginosa* virulence and intracellular survival. The diversity of gene functions and phenotypes affected by eDNA highlights the significant influence of this extracellular anionic polymer on the biology of *Pseudomonas aeruginosa*.

## Materials and Methods

### Strains, plasmids and media conditions

*P. aeruginosa* PAO1 was used as the wild-type strain and all the mini-Tn*5*-*lux*CDABE reporter strains listed in Table 1 (16). *P. aeruginosa* strains were routinely maintained at 37°C on Luria Broth (LB) agar plates and cultured in defined basal medium 2 (BM2) medium using succinate (20 mM) as the carbon source (10) and supplemented with extracellular fish sperm DNA (USB, 14405) and various magnesium concentrations, as indicated.

**Table 1.**
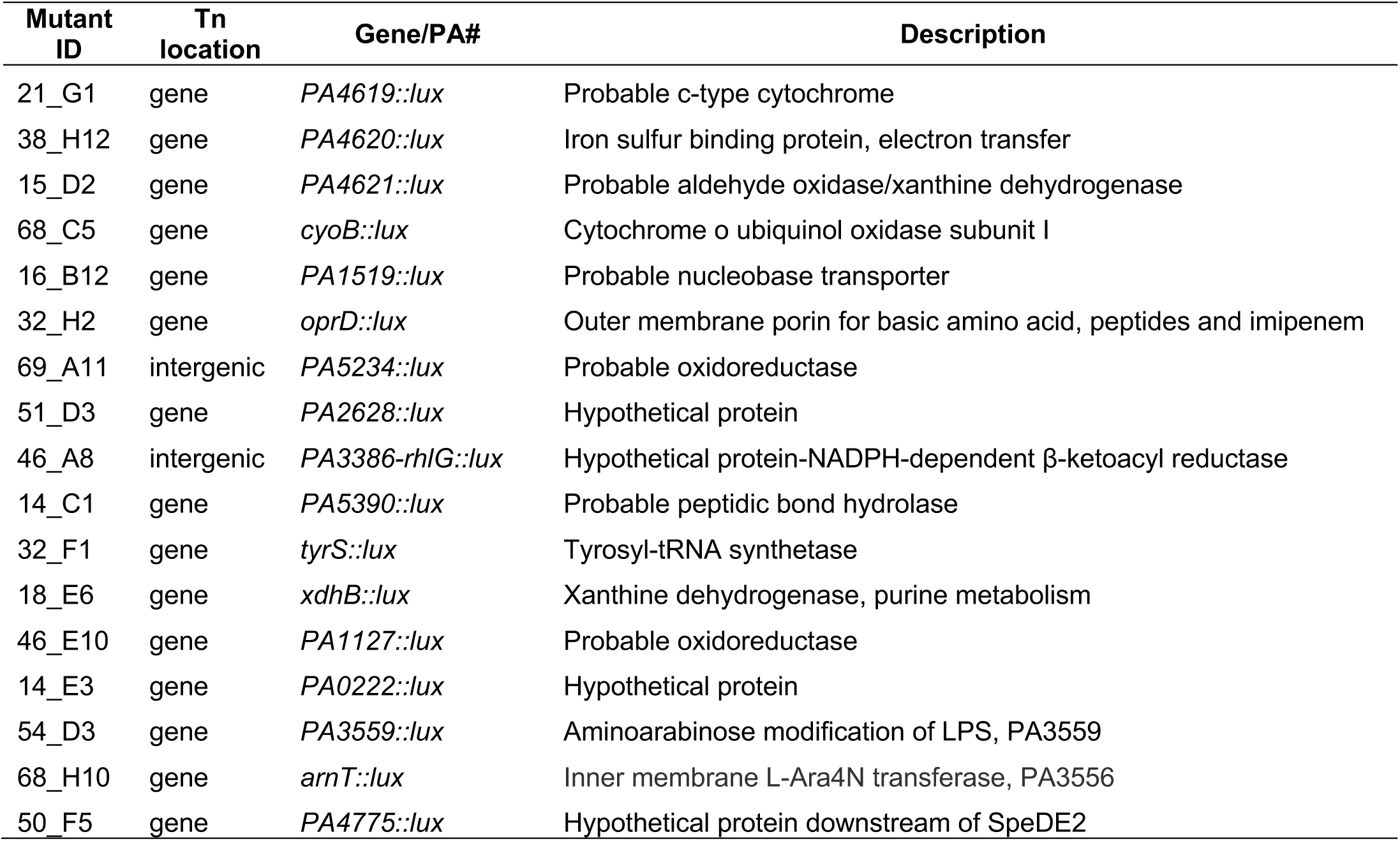
Transcriptional *luxCDABE* reporter strains used in this study.

| Mutant ID | Tn location | Gene/PA# | Description |
| --- | --- | --- | --- |
| 21_G1 | gene | *PA4619::lux* | Probable c-type cytochrome |
| 38_H12 | gene | *PA4620::lux* | Iron sulfur binding protein, electron transfer |
| 15_D2 | gene | *PA4621::lux* | Probable aldehyde oxidase/xanthine dehydrogenase |
| 68_C5 | gene | *cyoB::lux* | Cytochrome o ubiquinol oxidase subunit I |
| 16_B12 | gene | *PA1519::lux* | Probable nucleobase transporter |
| 32_H2 | gene | *oprD::lux* | Outer membrane porin for basic amino acid, peptides and imipenem |
| 69_A11 | intergenic | *PA5234::lux* | Probable oxidoreductase |
| 51_D3 | gene | *PA2628::lux* | Hypothetical protein |
| 46_A8 | intergenic | *PA3386-rhlG::lux* | Hypothetical protein-NADPH-dependent β-ketoacyl reductase |
| 14_C1 | gene | *PA5390::lux* | Probable peptidic bond hydrolase |
| 32_F1 | gene | *tyrS::lux* | Tyrosyl-tRNA synthetase |
| 18_E6 | gene | *xdhB::lux* | Xanthine dehydrogenase, purine metabolism |
| 46_E10 | gene | *PA1127::lux* | Probable oxidoreductase |
| 14_E3 | gene | *PA0222::lux* | Hypothetical protein |
| 54_D3 | gene | *PA3559::lux* | Aminoarabinose modification of LPS, PA3559 |
| 68_H10 | gene | *arnT::lux* | Inner membrane L-Ara4N transferase, PA3556 |
| 50_F5 | gene | *PA4775::lux* | Hypothetical protein downstream of SpeDE2 |

### Library construction, SOLiD sequencing and RNA-Seq analysis

For RNA isolation, *P. aeruginosa* PAO1 was grown in defined BM2 (2mM Mg^2+^) medium (10) as the negative control condition, and in BM2 (2mM Mg^2+^) medium supplemented with 0.75% (w/v; 7.5 mg/ml) extracellular DNA (USB, 14405). Total RNA was isolated from triplicate, mid-log cultures (3×10^9^ cells) using the RiboPure™-Bacteria Kit (Ambion) and treated with DNase (Ambion) until RNA was shown to be DNA-free using the Agilent Bioanalyzer. All subsequent steps were performed by the sequencing service company EdgeBio (Gaithersburg, MD). The Ribominus kit (Life Technologies) was used to remove rRNA from total RNA, and the cDNA library was constructed using the RNA-SEQ SOLiD kit. SOLiD 4 sequencing was performed on ∼150bp DNA fragments. Paired end reads were mapped to the PAO1 genome (17) using the RNA-SEQ Analysis Tool (CLCbio). In addition to manually removing rRNA sequences from the total reads, a de-duplication analysis using MarkDuplicates (Picard) was performed to remove duplicate sequences. Mapped reads were imported into CLCbio to determine the gene counts for each gene in each sample, which were used for differential gene expression analysis performed with DESeq (18). Genes were considered differentially expressed with fold changes greater than 2-fold and significance was determined by FDR values < 0.05.

### *lux* reporter gene expression assays

To validate eDNA-induced gene expression patterns, we selected a panel of transcriptional *luxCDABE* fusions from a library of mini-Tn*5*-*lux* mutants and performed gene expression assays in BM2 (16). While the transcriptome experiments were performed in standard BM2 + 0.75% (pH∼6), here used BM2 medium with and without supplementary 0.5% DNA (USB, 14405) and 0.1 mM Mg^2+^ at neutral pH 7. To neutralize the pH of BM2, we used 2X HEPES buffer (200 mM) to resist the pH changes associated with DNA addition. Standard BM2 containing 1mM Mg^2+^ was also adjusted to pH 5.5 and 7. Gene expression (CPS) was normalized to cell growth (optical density, OD_600_).

### Acid tolerance assays

*P. aeruginosa* PAO1 was grown overnight in LB, LB adjusted to pH 5.5 or LB supplemented with 0.2% DNA, then sub-cultured in the same corresponding conditions until mid-log. Cultures were then normalized to similar OD_600_ values and cells (5×10^7^ CFU) were resuspended in LB adjusted to pH 3.5 to subject the cells to acid shock. At various time points after acid shock (5-120 minutes), cells were serially diluted and plated to count the surviving population.

### Phagocytosis and intracellular survival experiments

RAW 264.7 cells (mouse leukemic monocyte macrophage cell line) were seeded at 5×10^5^ macrophages per well of a 24-well plate 24h prior to infection in (DMEM) with L-glutamine (2 mm) (Invitrogen). Overnight cultures of *P. aeruginosa* PAO1 or mutants were grown in relevant media were washed to remove extracellular components. Bacteria were in PBS pH 7.4 and bacterial densities adjusted so as to infect cells at a multiplicity of infection (MOI) of approximately 50:1. Phagocytosis was allowed to proceed for 1h, followed by a 4h incubation with 100 μg/ml of polymyxin B to kill extracellular bacteria (19). At 5h post-infection DMEM media was removed and infected macrophages treated for 15 mins with 0.1% Triton-X-100 (Sigma). Intracellular survival of *P. aeruginosa* 5h post-infection was quantified by serial dilution and plating of lysed macrophages on LB plates.

### Nicking *P. aeruginosa* infection of Drosophila

*Drosophila* were maintained routinely on medium containing corn meal, agar, sucrose, glucose, brewers’ yeast, living yeast, propionic acid, and phosphoric acid (20). Fly nicking assays were performed as previously described (21) using 3-5 day old female flies. Fly survival was monitored and recorded from 12 to 24 h post-inoculation. Kaplan-Meier survival curves were plotted, and statistical analysis was performed using GraphPad Prism 5 software. Significant differences in *Drosophila* survival were determined using the log-rank test.

### Inductively Coupled Plasma Mass Spectrometry (ICP-MS) for metal analysis

DNA solutions (10 mg/ml) were sent to Exova (Edmonton, Alberta) for ICP-MS analysis to quantitate a panel of metals bound to DNA. Three different DNA samples were submitted including fish DNA (USB, 14405), Na^+^-DNA (USB, 14377) and K^+^-DNA (USB, 14376).

### Funding Information

This research was supported by a Cystic Fibrosis Canada operating grant and the Westaim-ASRA Chair in Biofilm Research, both held by SL. HM was supported by a Cystic Fibrosis Canada postdoctoral fellowship.

## Results

### RNA-seq analysis to identify the transcriptome of eDNA-regulated genes

We performed RNA-seq analysis of *P. aeruginosa* PAO1 in the absence and presence of extracellular DNA (0.75%; 7.5 mg/ml) added to BM2 defined medium in order to identify the global profile of genes regulated by eDNA. Triplicate cultures with and without eDNA were grown to mid-logarithmic phase for total RNA isolation. DNA-free total RNA was depleted for rRNA and used to prepare cDNA libraries for SOLiD 4 sequencing. After mapping the initial sequence reads to the PAO1 genome, there were significant rRNA sequences remaining despite rRNA depletion, which were manually removed from the raw data. In addition, duplicate sequences from both conditions were identified and removed before performing DESeq analysis to identify differentially regulated genes.

We previously identified two operons that are induced by the addition of extracellular DNA: the *arn/pmr* operon (*PA3552-PA3559*) that is required for the addition of aminoarabinose to the lipid A moiety of lipopolysaccharide (LPS), and the spermidine synthesis genes *PA4773-PA4774* (1). We considered these two operons as internal positive controls given their DNA-induced expression patterns using *lux* reporter fusions (9–11) and quantitative RT-PCR (10). Although many of these genes were induced by at least 2-fold, not all genes within these operons were significantly upregulated by DESeq analysis (Table S1). Given the general observation that many gene clusters or operons showed trends of induction by eDNA, we included all 255 genes that were induced by 2-fold by the presence of extracellular DNA in Supplementary Table 1. In total, there were 148/255 genes that were induced by at least 2-fold (p<0.05), and 89/255 genes that were considered significantly induced by eDNA (FDR<0.05). We limit our discussion to mainly those genes that were significantly regulated by eDNA.

To summarize, there were several categories of genes induced by eDNA. The majority of genes appear to be required for basic metabolism and energy generation. This group includes numerous enzymes in basic metabolic enzymes and nutrient uptake pathways, such as nucleotide/nucleoside transport and catabolic genes. This group is likely required in the utilization of DNA as a nutrient source. There were also categories of genes that could be involved in antibiotic resistance, metal transport/efflux, and numerous transcriptional regulators (Table S1). In addition to genes that were induced, there were also genes that were significantly repressed by the addition of extracellular DNA. In total, there were 109 genes that were repressed by at least 2-fold (p<0.05), and 76/109 genes that were considered significantly repressed by eDNA (FDR<0.05). The major categories of DNA-repressed genes are involved in metabolism, metal efflux, and bacterial secretion systems (Table S2).

### DNA-induced metabolic genes are likely required to utilize DNA as a nutrient

The majority of eDNA-induced genes appear to be involved in various aspects of central metabolism (Table S1). The most intuitive induced genes were those annotated with functions in nucleotide/nucleoside metabolism or transport. For example, the xanthine dehydrogenase cluster (*xdhABCD*) is a complex molybdenum-containing flavoprotein that is involved in purine catabolism and may be involved in electron transport processes. Within the same cluster, the guanine deaminase *PA1521* and the *PA1517-PA1513* operon are also likely involved in purine catabolism (Table S1). The cytosine deaminase *PA0142* may contribute to pyrimidine catabolism. Various porins (*opdC, opdH, oprD*) and transporters (*PA2938, PA0030, PA0222, PA0273, PA3079, PA4355, PA4622, PA4654, PA0220, PA5158*), are induced by eDNA, which may be involved in the uptake of DNA, short oligonucleotides, nucleotides or polyamines (Table S1). There is a large group of DNA-induced genes annotated for diverse biosynthetic and catabolic enzymes, and energy generation processes (Table S1). For example, the *cyoABCDE* genes encode a cytochrome o ubiquinol oxidase subunit II (22) and may play a role in electron transport and energy generation in the presence of eDNA (Table S1).

It is also interesting that specific metabolic genes are also repressed by extracellular DNA (Table S2). The Cbb3-type cytochrome oxidase (PA4133) gene is repressed and supports the concept that environmental conditions influence the expression of the aerobic, terminal oxidases of the electron transport process (22). The *PA3519-PA3515*, cluster contains a probable adenylosuccinate lyase involved in purine biosynthesis (PA3517, PA3516). While nucleotide catabolism genes are induced, the biosynthetic genes can be repressed, when in the presence of DNA as a nutrient source.

### Antibiotic resistance genes are induced by extracellular DNA

We have previously shown that spermidine synthesis genes *PA4773-PA4775* and the aminoarabinose modification of lipid A genes *PA3552-PA3559* (*arn/pmr*) are two operons required for DNA-induced antimicrobial peptide resistance (1). The modifications protect the outer membrane from cationic antimicrobial peptide damage in DNA-enriched biofilms or planktonic cultures (9,10), and limit aminoglycoside permeability under acidic conditions (11). They also protect the outer membrane from direct DNA damage and killing by neutrophil extracellular traps (3). These genes are highly expressed in the presence of eDNA and are required for shielding the outer membrane and protecting from diverse membrane and antibiotic threats.

The transcriptome of eDNA-induced genes included other potential antibiotic resistance genes such as a β-lactamase (*PA2315*), an aminoglycoside phosphotransferase (*PA1829*), and the OpmG outer membrane protein that contributes to aminoglycoside resistance (23), which is adjacent to the *emrAB* multidrug efflux pump (*PA5157-PA5159*) (Table S1). Interestingly, eDNA represses the expression of *nalD* (Table S2), which is a transcriptional repressor of the MexAB-OprM RND efflux pump (24). This regulatory effect may ultimately lead to activation of the MexAB-OprM pump in the presence of eDNA. These systems may act as novel resistance determinants that are uniquely expressed in DNA rich biofilms or infection sites.

### DNA influences the expression of diverse metal efflux and transport pathways

An interesting subset of the transcriptome included genes that are involved in both the efflux and transport of metal cations. Among the DNA-induced genes, there are numerous transport systems for sodium (Na^+^), iron (Fe^2+^) and zinc (Zn^2+^) (Table 2), mostly involved in the uptake of these cations. The TerC protein is an integral membrane protein that effluxes tellurium and therefore contributes to tellurium resistance (Table 2). Interestingly, among the DNA-repressed genes, there are numerous RND efflux and transport systems for cobalt (Co^2+^), zinc (Zn^2+^) and cadmium (Cd^2+^). The repression of metal efflux pumps and induction of cation transport pathways could collectively lead to increased rates of metal cation uptake (Table 2).

**Table 2.** Extracellular DNA-repressed and induced metal transport, efflux and regulatory systems.

| **p value** | **FDR value** | **Fold change** | **Gene ID** | **Description of Repressed Gene** |
| --- | --- | --- | --- | --- |
| 1.1329E-24 | 4.8741E-22 | 49.17 | PA3520 | Probable copper chaperone |
| 1.25495E-78 | 3.5164E-75 | 33.85 | PA3920 | Metal transporting P-type ATPase |
| 9.32E-43 | 1.7411E-39 | 31.28 | PA2521 | RND metal cation efflux membrane protein CzcB |
| 1.02606E-12 | 1.85E-10 | 30.91 | PA3522 | RND efflux transporter |
| 5.61431E-40 | 7.8628E-37 | 30.57 | PA3523 | RND efflux membrane fusion protein |
| 1.11E-26 | 5.1921E-24 | 29.71 | PA2522 | Outer membrane protein CzcC |
| 2.32503E-31 | 1.8597E-28 | 22.50 | PA2523 | Two-component response regulator CzcR |
| 1.14965E-39 | 1.2878E-36 | 20.29 | PA2520 | RND divalent metal cation efflux transporter CzcA |
| 7.54787E-12 | 1.2014E-09 | 12.13 | PA2524 | Two-component sensor CzcS |
| 7.20894E-13 | 1.3864E-10 | 10.05 | PA3690 | Metal-transporting P-type ATPase |
| 0.000312263 | 0.01198629 | 8.50 | PA3521 | Probable outer membrane protein |
| 7.63089E-05 | 0.00347113 | 5.25 | PA2807 | Hypothetical protein |
| 0.000625388 | 0.02161567 | 4.42 | PA0397 | Co<sup>2+</sup>, Zn<sup>2+</sup>, Cd<sup>2+</sup> efflux system protein |
| 0.001934937 | 0.05975273 | 3.06 | PA1297 | Metal cation transporter |
| **p value** | **FDR value** | **Fold change** | **Gene ID** | **Description of Induced Gene** |
| 1.22308E-09 | 1.4466E-07 | 4.56 | PA2026 | Probable Na<sup>+</sup>-dependent transporter |
| 1.55153E-05 | 0.00082963 | 2.64 | PA3790 | Copper transport outer membrane porin OprC |
| 1.79203E-05 | 0.00094895 | 2.54 | PA2549 | TerC membrane protein |
| 0.002904417 | 0.08727124 | 2.52 | PA3749 | Major facilitator superfamily (MFS) transporter |
| 0.000444716 | 0.0163311 | 2.19 | PA5248 | High-affinity Fe<sup>2+</sup>, Pb<sup>2+</sup> permease |

We have shown previously that DNA can efficiently bind to exogenous divalent metal cations such Mg^2+^, Ca^2+^, Mn^2+^ and Zn^2+^ (9). To confirm that DNA sequesters diverse metals, we analyzed various commercial DNA preparations for total bound metals using inductively coupled plasma mass spectrometry (ICP-MS). All DNA samples bound to a wide range of metals. As internal controls, we confirmed that that Na^+^-DNA bound the highest amount of Na^+^, and the K^+^-DNA bound the highest amount of K^+^ (Table 3). These samples were likely named Na^+^-DNA or K^+^-DNA as a consequence of DNA precipitation with these respective salts during the purification process. The highly bound cations and metals to DNA included Na+, K+, Ca^2+^, Mg^2+^, Zn^2+^, Cr^3+^, Sr^2+^, Cd^2+^, Co^2+^, Fe^2+^, Pb^2+^, Al^3+^ and Mn^2+^ (Table 3).

**Table 3.** Metals bound to commercially available sources of purified DNA.

| Metal <sup>a</sup> | DNA | Na<sup>+</sup> DNA | K<sup>+</sup> DNA | Detection Limit |
| --- | --- | --- | --- | --- |
|  | mg/L |  |  | mg/L |
| Sodium | 158 | 583 | 5.4 | 0.4 |
| Sulfur | 15 | 2.1 | 2.8 | 0.3 |
| Potassium | 3 | 2.4 | 817 | 0.4 |
| Calcium | 2 | 2.1 | 2 | 0.2 |
| Magnesium | 1.7 | 0.93 | 3.72 | 0.2 |
| Zinc | 0.1 | 0.15 | 0.236 | 0.001 |
| Manganese | <0.02 | <0.01 | <0.01 | 0.005 |
| Vanadium | 0.035 | 0.069 | 0.0053 | 0.0001 |
| Boron | 0.03 | 0.005 | <0.004 | 0.002 |
| Nickel | 0.0078 | 0.0096 | 0.001 | 0.0005 |
| Chromium | 0.0059 | 0.0229 | 0.0044 | 0.0005 |
| Copper | 0.005 | 0.01 | 0.009 | 0.001 |
| Strontium | 0.005 | 0.01 | 0.005 | 0.001 |
| Lead | 0.0006 | 0.0004 | 0.001 | 0.0001 |
| Cadmium | 0.00011 | 0.00002 | 0.00002 | 0.00001 |
| Iron | <0.3 | 0.58 | 0.1 | 0.05 |
| Molybdenum | <0.005 | 0.008 | <0.002 | 0.001 |
| Barium | <0.005 | 0.003 | 0.003 | 0.001 |
| Selenium | <0.001 | 0.0007 | 0.0007 | 0.0002 |
| Cobalt | <0.0005 | 0.0002 | <0.0002 | 0.0001 |

<sup>a</sup>The following metals were detected at very low levels, near or below the limit of detection: silver, thallium, lithium, tin, zirconium, antimony, arsenic, bismuth, titanium and uranium.

The CzcRS two-component system activates expression of the CzcCBA efflux system in the presence Co^2+^, Cd^2+^ and Zn^2+^ (25,26). DNA binds and sequester Co^2+^, Cd^2+^ and Zn^2+^ (Table 3), thereby limiting exposure to these metals, which may explain why the the CzcRS regulatory system and the CzcCBA efflux system are repressed by extracellular DNA (Table S2). These metals are required for growth in trace amounts and are toxic in higher concentrations, and therefore require homeostatic uptake processes to balance the need and toxicity of these metals. In conclusion, we propose that the anionic phosphate backbone of DNA permits the binding and sequestering of diverse metal cations, which leads to a complex response to achieve a balance in metal uptake.

### Extracellular DNA repressed the H2-T6SS and the T3SS

The type III secretion system (T3SS) is required to deliver exotoxins directly into host cells through a needle-like structure, which interfere with host cell responses and contribute to bacterial virulence (27). The type VI secretion system (T6SS) encodes a contractile syringe structure required for interbacterial killing and allows for competition among mixed bacterial communities (28). Extracellular DNA repressed the expression of many genes within the T3SS and the H2-T6SS clusters (Table S2). While DNA specifically represses the H2-T6SS (Table S2), previous studies have demonstrated that cation chelation by eDNA can rapidly activate the H1-T6SS through post-translational control, which leads to nonselective attack and killing of neighbouring bacterial species (29). The H1-T6SS contributes primarily to bacterial killing, but the H2-T6SS contributes to killing of eukaryotic and bacterial cells (28).

The type III secretion system is controlled by limiting Ca^2+^, and we have shown that various, exogenous Ca^2+^ chelators, such as NTA, alginate and DNA, can induce the expression of the T3SS (30). Here we report that using higher DNA concentrations from a different source than previously tested (30) represses the T3SS. Therefore, it is likely that DNA can have various effects on the T3SS based on the relative concentrations and chelation potential. While limiting Ca^2+^ functions as the inducing signal, limiting both Mg^2+^ and Ca^2+^ cations represses the T3SS (30). These results are ultimately consistent with the role of cations concentrations in controlling the T3SS and T6SS.

### Extracellular DNA influences gene expression through cation chelation, acidification or as a nutrient

In order to validate the novel eDNA-regulated genes identified by RNA-seq analysis, we searched our mini-Tn*5*-*lux* transposon mutant library (16) for transcriptional *lux* reporters to genes in Supplementary Table 1 (Table 1). There are multiple ways in which eDNA can influence gene expression. DNA is a cation chelator and therefore creates a Mg^2+^ limiting condition and induces genes that are known to be controlled by magnesium limitation and the PhoPQ/PmrAB two component systems (9,10). The accumulation of eDNA acidifies biofilm cultures and acidic pH triggers expression of the PhoPQ PmrAB-controlled genes (11). To separate the effects of cation chelation and pH, we measured the expression of candidate DNA-induced genes under: 1) eDNA and neutral pH 7 and 2) acidic pH 5.5 or pH 7 with excess 1 mM Mg^2+^, in order to provide insight as to how DNA influences the global gene expression phenotypes of *P. aeruginosa*.

The first objective was to validate the gene expression responses to eDNA observed in the transcriptome. Transcriptional fusions to known DNA-induced genes *PA4775::lux* and *PA3559::lux* were used as positive controls to show induction by eDNA at neutral pH (Fig 1). The baseline expression of these reporters is high in baseline conditions without DNA, since 100 μM Mg^2+^ is already an inducing condition, and eDNA induces expression further by 2-fold (Fig 1). However, many genes have very low expression levels in the absence of DNA and are induced very strongly. For example, several metabolic genes showed a strong induction response to eDNA, including *cyoB, oprD, xdhB, PA1519, PA4620, PA4621, tyrS* and *PA5234* (Fig 1). In comparison, some genes had low levels of baseline expression and showed modest induction by eDNA (PA1127, PA0222) (Fig 1)

**Fig 1.**
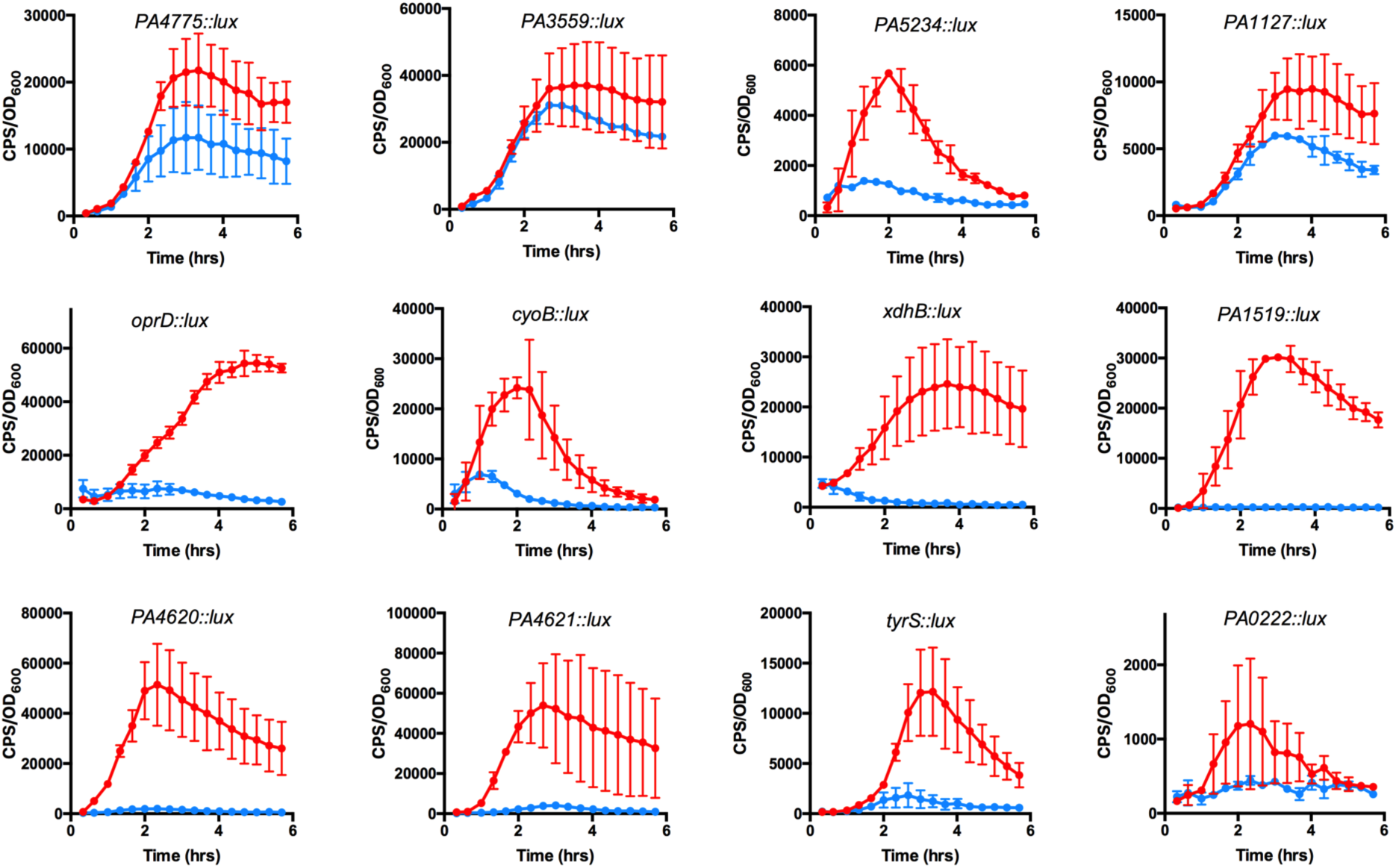
Extracellular DNA-induced gene expression patterns. Transcriptional *lux* fusion reporter strains were grown in BM2 medium containing 100 μM Mg^2+^ (pH 7) without (blue) or with the addition of 0.5% DNA (red) and gene expression was measured every 20 min throughout 6 hrs. Values shown are the averages +/- SEM from duplicate values, and each experiment was repeated 3-6 times. Each panel is labelled with the DNA-induced gene of interest.

Since it is known that acidification by eDNA is a separate signal to induce expression of the *arn/pmr* and *PA4773-PA4774* operons, we wanted to determine if any of the novel eDNA-induced genes were responding to acidic pH. We therefore measured the expression of these reporters in neutral pH 7 and under mild acid pH 5.5. Figure 2 demonstrates that some eDNA genes are strongly induced by pH 5.5, including the *oprD* outer membrane porin and the Cyo cytochome oxidase, and others are modestly induced by acid pH (PA2628). Both *oprD* and *cyoB* are not regulated by limiting Mg^2+^ (data not shown), and therefore respond to DNA as a possible nutrient source or to pH changes associated with the addition of eDNA. Induction of these genes by acidic pH suggests a possible role in adjusting to the stress of acidic pH. To summarize, we validated the DNA-induced gene expression phenotypes from the transcriptome using transcriptional *lux* fusions and illustrated the possible mechanisms for DNA influencing gene expression through cation chelation, acidification or as a nutrient source.

**Fig 2.**
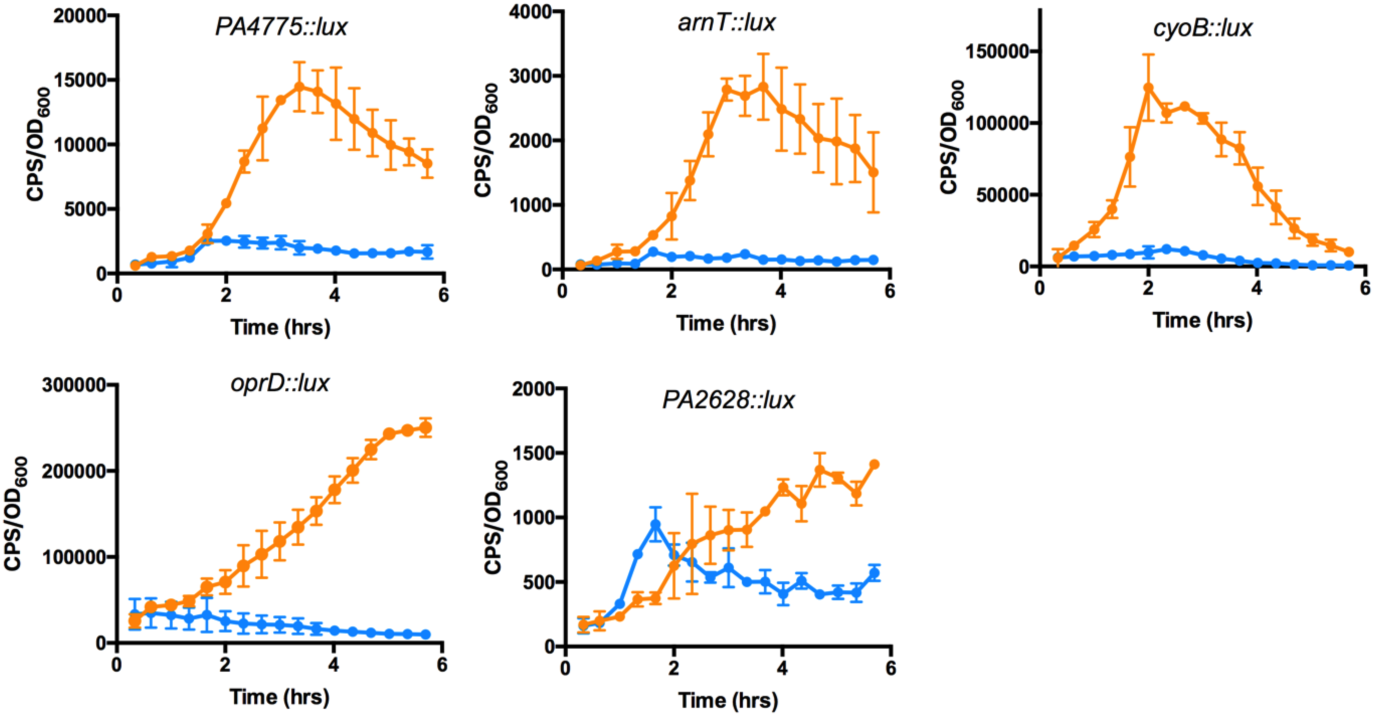
pH-induced gene expression patterns. Transcriptional *lux* fusion reporter strains were grown in BM2 medium containing 1mM Mg^2+^ at pH 5.5 (orange) or pH 7 (blue) and gene expression was measured every 20 min throughout 6 hrs. Values shown are the averages +/- SEM from duplicate values, and each experiment was repeated 3-6 times. Each panel is labelled with the DNA-induced gene of interest.

### Extracellular DNA promotes an acid pH tolerance response

*Salmonella typhimurium* is known to have an acid tolerance response (ATR) that may contribute to survival during passage through the stomach and gastrointestinal tract or within macrophages (31). Since *P. aeruginosa* induces specific genes during mild acid exposure, we sought to determine if *P. aeruginosa* has an acid tolerance response, and if growth in the presence of eDNA promoted survival upon challenge with a lethal pH 3.5 acid shock. PAO1 cultures were grown overnight in LB, LB supplemented with 0.2% DNA, or LB adjusted to pH 5.5. Overnight cultures were sub-cultured and grown to mid-log phase in the same corresponding conditions, before being subjected to an acid shock (pH 3.5). Colony counts were performed before and after the acid shock to determine the relative survival of the three populations. After 20 to 60 minutes exposure to acid pH, the cultures grown in mild acid pH 5.5 were shown to promote an acid tolerance response. Interestingly, growth in presence of 0.2 % extracellular DNA led to the greatest survival and tolerance to exposure to pH 3.5 shock treatment. After 90 minutes of exposure to acid pH 3.5, LB grown cultures decreased to zero viability. However, PAO1 that was pre-grown in extracellular DNA was the only condition that promoted survival for up to 120 minutes (Fig 3). We recently demonstrated that DNA accumulation results in acidic microdomains in biofilms and pH values decreased to ∼5.5 in biofilms formed by an eDNA hyperproducing strain (11). Given the ubiquitous accumulation of eDNA in biofilms, we propose that the acid tolerance response is required to survive the pH gradients that establish within biofilms.

**Fig 3.**
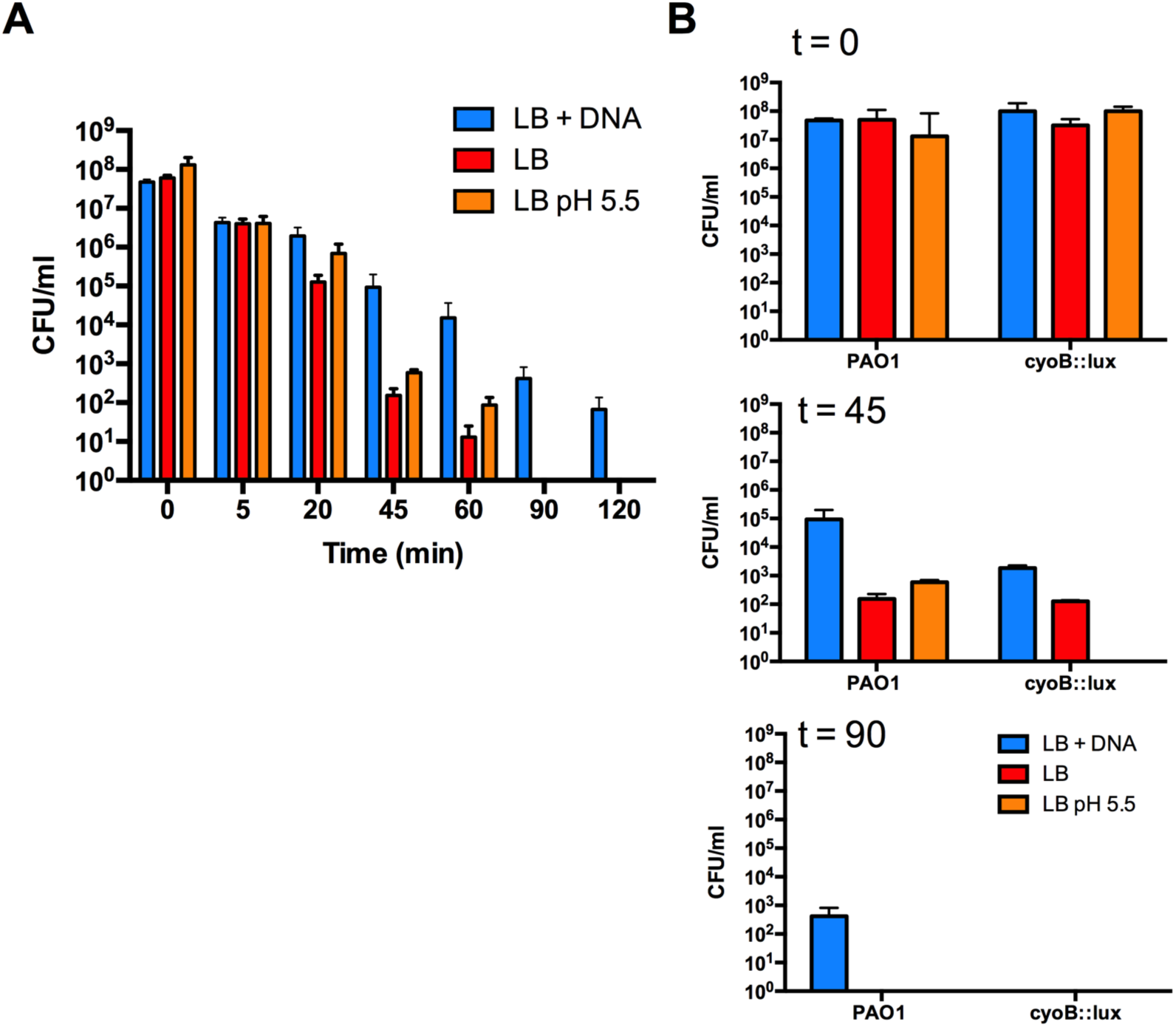
Extracellular DNA promotes an acid tolerance response in *P. aeruginosa*. (A) Wild type PAO1 was grown overnight in LB, LB containing 0.2% DNA or LB adjusted to pH 5.5, and then exposed to an acid shock of pH 3.5. The cultures were plated to enumerate bacterial survival at 15 min intervals after the acid shock. Values shown represent the average and standard error of triplicate CFU/ml values, and each experiment was performed three times. (B) Wild type PAO1 and the *cyoB::lux* mutant were grown overnight in LB, LB containing 0.2% DNA or LB adjusted to pH 5.5, and then exposed to an acid shock of pH 3.5. Bacterial numbers were determined before the acid shock (t=0) and then at 45 and 90 min after being exposed to the pH 3.5 acid shock. Values shown represent the average and standard error of triplicate CFU/ml values, and each experiment was performed three times.

To further understand the possible mechanisms of acid tolerance, we hypothesized that specific acid pH-induced genes may contribute to acid tolerance. Two common pH homeostasis mechanisms under acidic conditions are the transport or efflux of protons out of the cytoplasm, and the consumption of protons through metabolic reactions (32). Proton transport mechanisms include H^+^ exchange proteins, Na^+^/H^+^ antiporters, and the electron transport process (32). Proton consumption occurs through the expression of hydrogenases and decarboxylases that consume protons during their reactions (32).

*Pseudomonas aeruginosa* encodes five aerobic, terminal oxidases that are differentially regulated dependent on the growth conditions, and operate as low or high oxygen affinity systems during the membrane-bound electron transport process (22). The *cyoABCDE* quinol oxidase is a low affinity system and would be expected to operate when oxygen is abundant during exponential growth, although the cyo genes are weakly expressed under normal growth conditions in Luria Broth (22). Strong induction of the *cyoABCDE* genes under acidic pH suggests that this oxidase is efficient for proton transport under acidic conditions (Fig 3). To confirm the role of the *cyo* quinol oxidase in surviving an acid pH shock, we compared the acid tolerance response of the wild type strain to a transposon mutant in the *cyoB::lux* gene. The *cyoB::lux* mutant did not survive an acid shock treatment beyond 45 minutes when pre-grown in mild acid pH 5.0, and did not survive beyond 90 min when-grown or in the presence of 0.2% eDNA, respectively (Fig 3). These data suggest that the *cyoABCDE* genes are required to balance cytoplasmic pH during an acid shock, and therefore contribute to the acid tolerance response.

### Growth in eDNA influences intracellular survival

We wanted to determine the influence of extracellular DNA on virulence phenotypes of *P. aeruginosa*. We hypothesized that DNA-induced antimicrobial peptide resistance (1), in combination with DNA-induced acid tolerance (Fig 3), would promote intracellular survival during phagocytosis. *P. aeruginosa* PAO1 was cultured overnight in BM2 supplemented with or without 0.75% DNA and bacterial densities were adjusted to infect fully confluent macrophage cell monolayers at a MOI of 50:1. A significant increase was observed in the number of CFUs recovered following 5 hrs of intracellular survival for PAO1 precultured in the presence of extracellular 0.75% DNA, and also when pre-cultured in limiting Mg^2+^ (20 μM). (Fig 4A). PAO1 grown in the presence of extracellular DNA and excess 10mM Mg^2+^ were not significantly differently to PAO1 precultured in BM2 with excess 2mM Mg^2+^ (Fig 4A). The addition of excess Mg^2+^ reduced the survival phenotype to wild type levels, presumably because exogenous magnesium neutralized the cation chelating effects of DNA (9).

**Fig 4.**
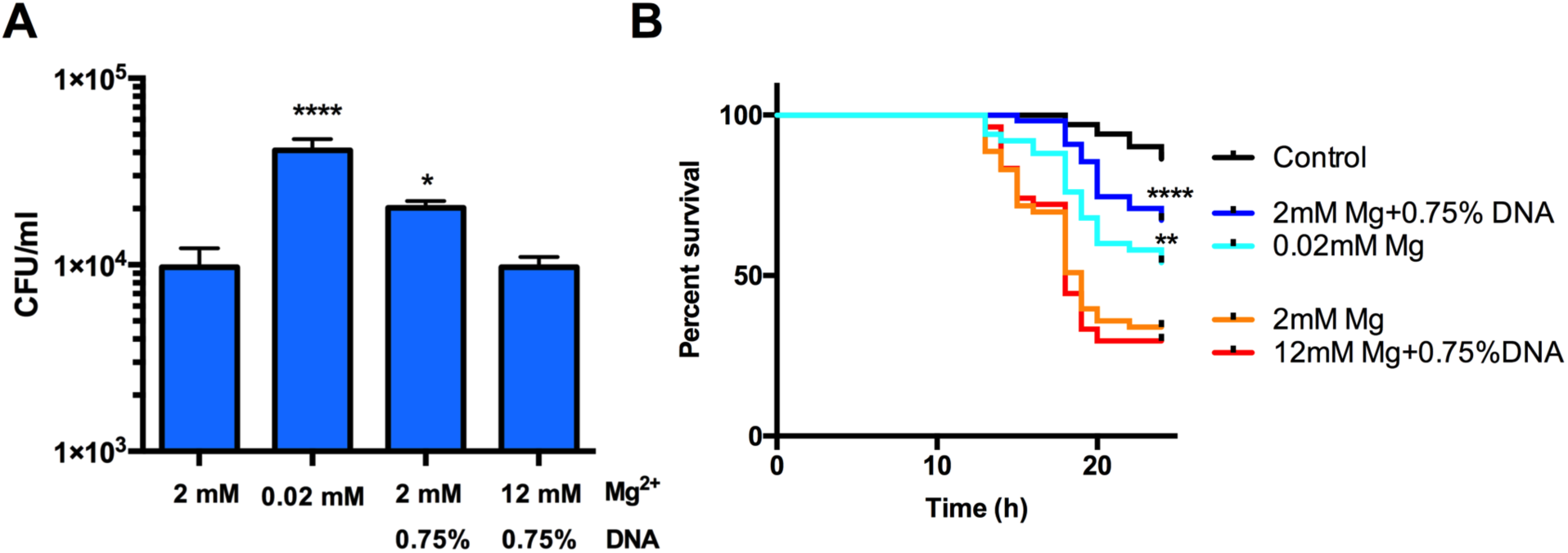
Extracellular DNA promotes intracellular survival during phagocytic killing and affects virulence of *P. aeruginosa* in a fly nicking model of infection. (A) Bacteria were recovered following 5h incubation with macrophages. Bacteria were precultured in BM2 2mM, 0.02mM Mg^2+^, BM2 2 mM Mg^2+^ + 0.75% DNA or 12 mM Mg^2+^ + 0.75% DNA (excess Mg) and macrophages were infected at an MOI of 50:1. Data is represented as the mean+/-SEM from three independent experiments. Bacteria grown in 0.02 mM Mg^2+^ and in 2 mM Mg^2+^ + 0.75% DNA showed significantly greater survival than cultures grown in 2mM Mg^2+^ (****p<0.0001; *p<0.05) as determined by one-way ANOVA with Bonferroni post-tests. (B) Kaplan-Meier survival curves of *Drosophila* post nicking infection with PAO1 grown in BM2 2mM, 0.02 mM Mg^2+^, 2mM Mg^2+^ + 0.75% DNA or 12mM Mg^2+^ + 0.75% DNA (excess Mg^2+^). Experiments were performed at least 3 times each with a minimum of 50 flies and representative curves are shown. Significant differences were determined with the log rank test between infections with bacteria grown in 0.02 mM and 2mM Mg^2+^ (**p<0.01), between 2mM Mg^2+^ and 2mM Mg^2+^ + 0.75% DNA (**** p<0.0001), as well as between 2 mM Mg^2+^ + 0.75% DNA and 12mM Mg^2+^ + 0.75% DNA (**** p<0.0001).

### Grown in eDNA influences virulence during fruit fly infections

The *Drosophila* nicking infection model (21) was used to assess the influence of eDNA on the virulence of *P. aeruginosa*. This infection model was preferred over the fly feeding model for this experiment, since fly killing is rapid in the nicking model (hours), and therefore the potential influence of the pre-growth conditions may have an immediate effect on the infection. In contrast, the slow killing kinetics of feeding infections would be less influenced by the phenotype of the ingested bacteria. *P. aeruginosa* PAO1 was grown in different conditions and injected into the abdomen of fruit flies, which results in fly death with 15 hrs. PAO1 grown in limiting Mg^2+^ or in 0.75% DNA were significantly less virulent (54-67% survival at 24h post-infection) than PAO1 pre-cultured in BM2 high Mg^2+^ or BM2 excess Mg^2+^ + 0.75% DNA (30-34% survival at 24h post-infection) (Fig 4B). The reduced virulence of PAO1 grown in the presence of eDNA may be related to the repression of the type III and type VI (H2) secretion systems (Table S2). These data indicate that *P. aeruginosa* pre-grown under conditions of cation limitation have altered virulence properties compared to strains grown in cation rich environments.

## Discussion

While extracellular DNA was initially discovered to have a structural role in maintaining the biofilm structure of young *P. aeruginosa* biofilms (33), we now understand that eDNA has many additional, non-structural functions. DNA imposes various stresses on cells by chelating and limiting the availability of metal cations, disrupting membrane integrity, acidifying the environment and acting as a nutrient (9,11). To gain further insight into how eDNA influences *P. aeruginosa*, we performed a transcriptome analysis. Correspondingly, many genes are induced in response to these stresses that contribute to defence and ultimately to long-term survival of *P. aeruginosa*.

In addition to the PhoPQ/PmrAB-controlled outer membrane modifications that protect against antimicrobial peptides, aminoglycosides, DNA and NET killing (16,9–11), eDNA induces the expression of additional antibiotic resistance genes. Extracellular DNA chelates metals and controls the expression of diverse metal uptake and efflux systems, in an attempt to acquire the limiting metals. The largest category of eDNA-induced genes are likely required to use DNA as nutrient source of phosphate, nitrogen or carbon. We demonstrate that similar to *S. typhimurium* (31), *P. aeruginosa* also has an acid tolerance response when grown with eDNA, which promotes survival to acid shock. The ability of *P. aeruginosa* to survive in acidic conditions is important in DNA-rich biofilms or infections sites, but also when ingested in the stomach, or when phagocytosed into acidified vacuoles.

Previous studies reported that magnesium limitation and PhoP repressed the expression of the *retS* orphan sensor that functions as a repressor of biofilm formation (34). Therefore, under magnesium limiting conditions, or in the presence of eDNA, *P. aeruginosa* produces increased amounts of the Pel/Psl exopolysaccharide, leading to robust aggregates and biofilms (34). Interestingly, we show here eDNA can repress the T3SS and the H2-T6SS, secretion systems required for virulence and protection from eukaryotic immune cells. Collectively, these eDNA-induced phenotypes are consistent with the proposed shift that controls acute and chronic infection phenotypes, and involves planktonic or biofilms modes of growth, respectively (35). Growth with eDNA reduced *P. aeruginosa* virulence during the acute, pin pricking infection model in *Drosophila*, which may be related to repression of the T3SS and T6SS.

Extracellular DNA promotes aggregation, antibiotic resistance and immune cell evasion phenotypes, which are the hallmark features of biofilms. This study expands our understanding of the significant influence of extracellular DNA on the physiology and gene expression patterns of *P. aeruginosa*. Although our experiments were performed in planktonic cultures containing eDNA, we propose that these results are relevant to biofilms and all conditions or infections where there is an accumulation of eDNA.

